## Supplementary figures for "Optogenetic frequency scrambling of hippocampal theta oscillations dissociates working memory retrieval from hippocampal spatiotemporal codes"

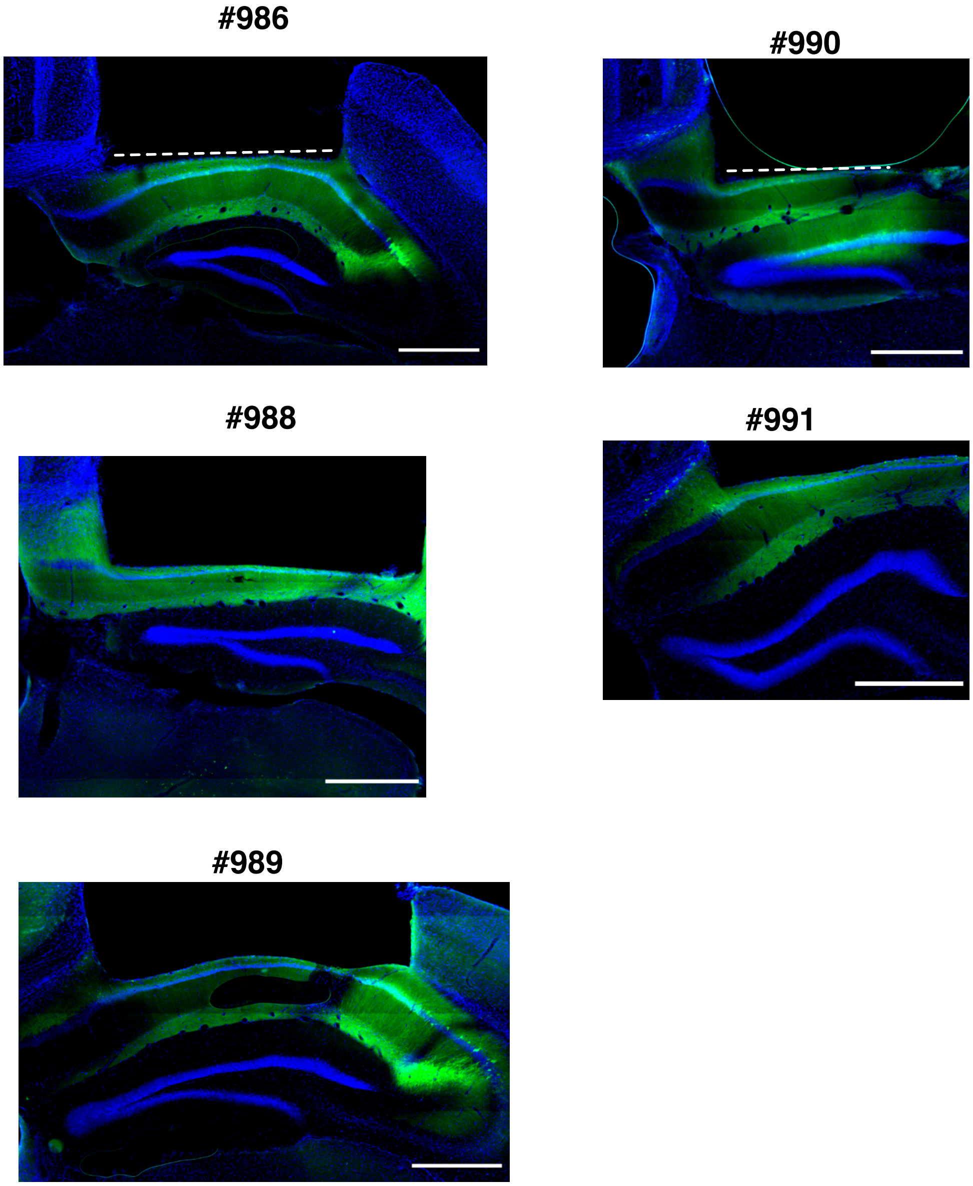


**Supplemental figure 1. Histological analysis of CKII-GCamp6f transfection and lens placement.** coronal sections of dorsal hippocampus displaying location of lens surface (white dotted line). blue, DAPI; green, GCamp6f. Scale bars in all micrographs: 500 µm.

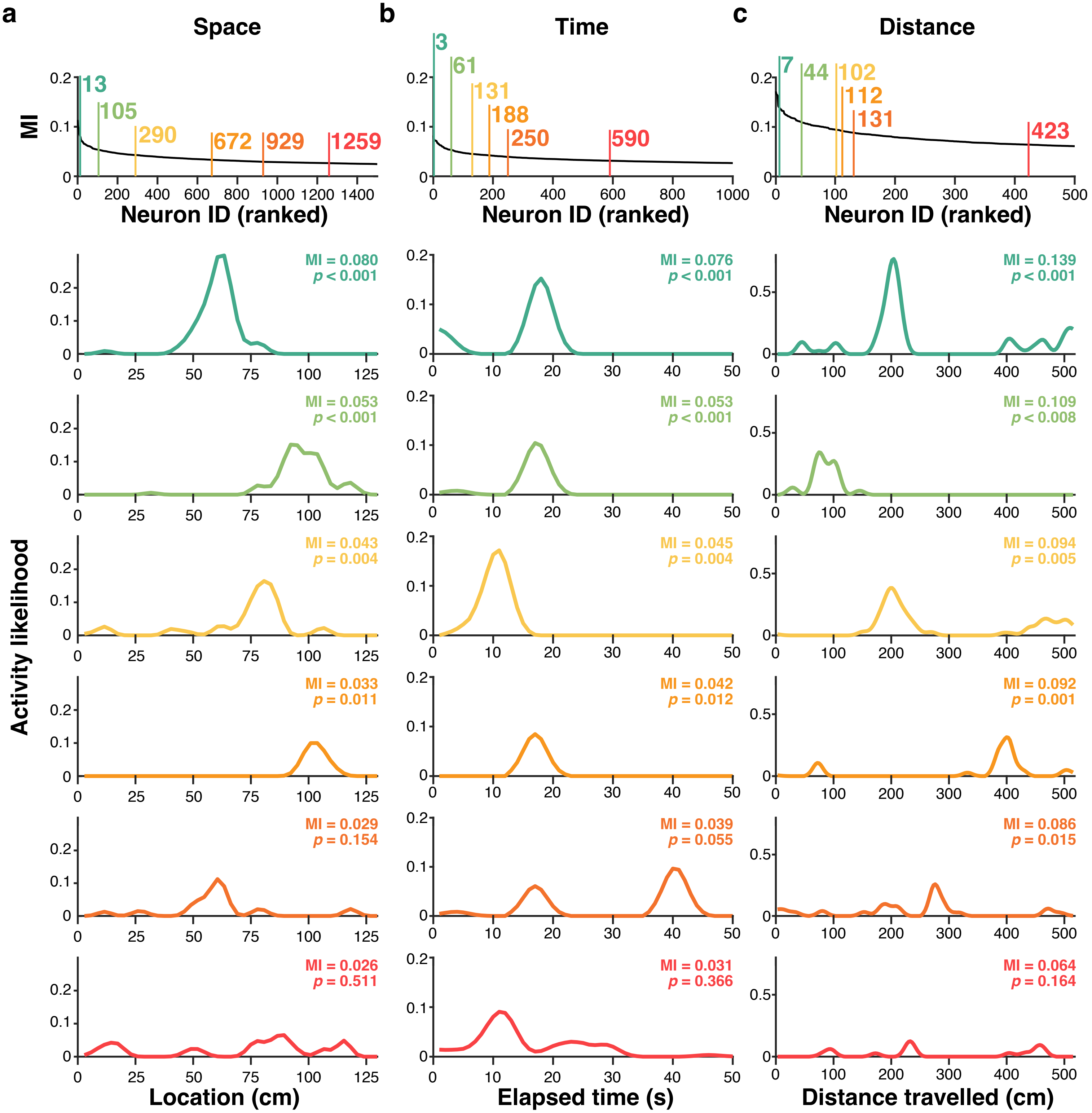


**Supplemental figure 2.** Example cells modulated by space (**a**), time (**b**), and distance travelled (**c**) and ranked by their mutual information. For each cell, a tuning curve with corresponding MI value and its significance (p-value) is displayed.

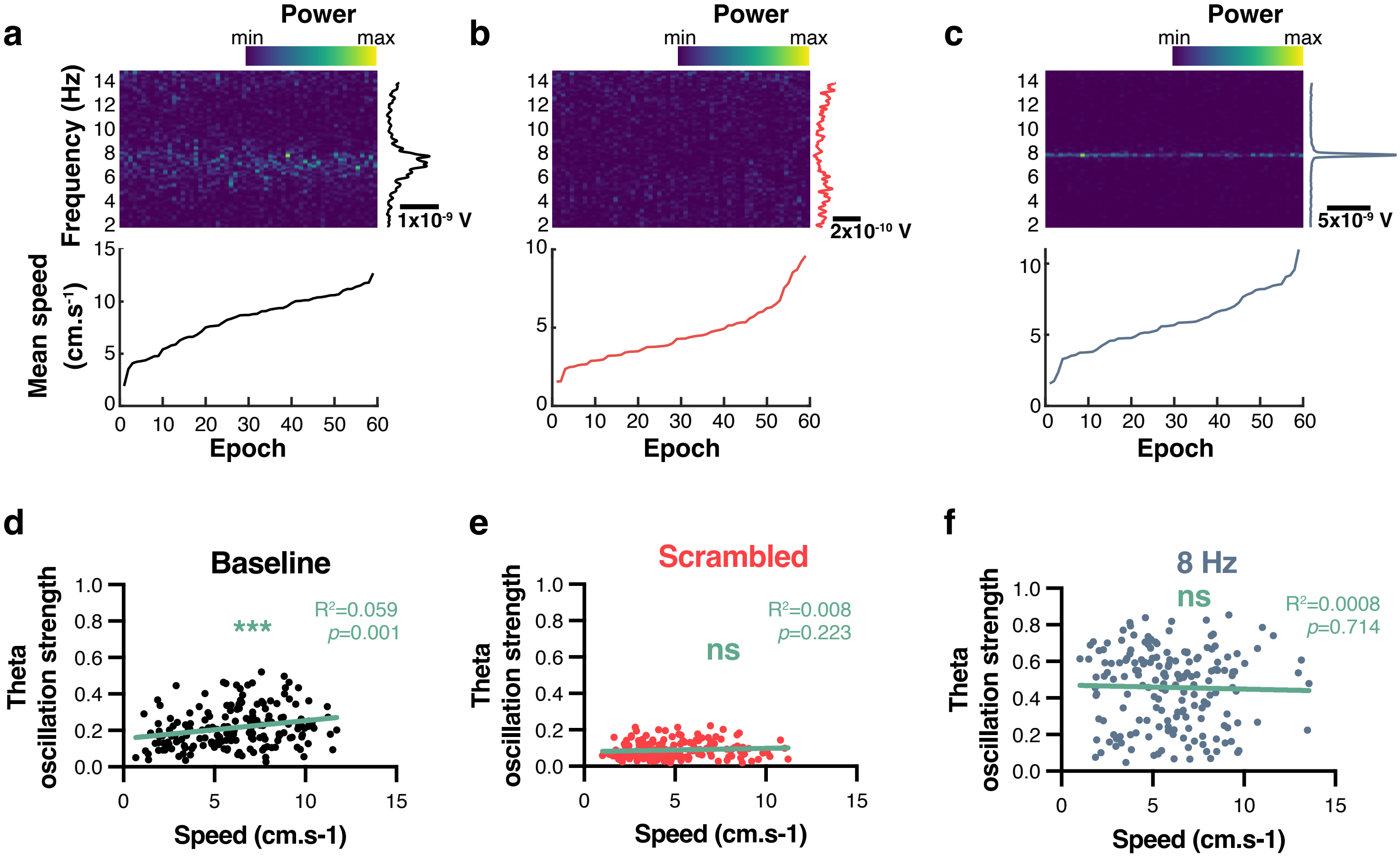


**Supplemental figure 3. Optogenetic stimulations control theta irrespective of locomotor speed. a**, baseline epochs containing natural theta. Top, spectrogram with x-axis sorted by average speed during recorded epoch. Bottom, corresponding locomotor speed. **b**, same for epochs during scrambled stimulations. **c**, same for epochs durings 8 Hz stimulations. **d**, **e**, **f**, theta oscillation strength against locomotor speed for baseline, scrambled, and 8 Hz stimulations respectively.

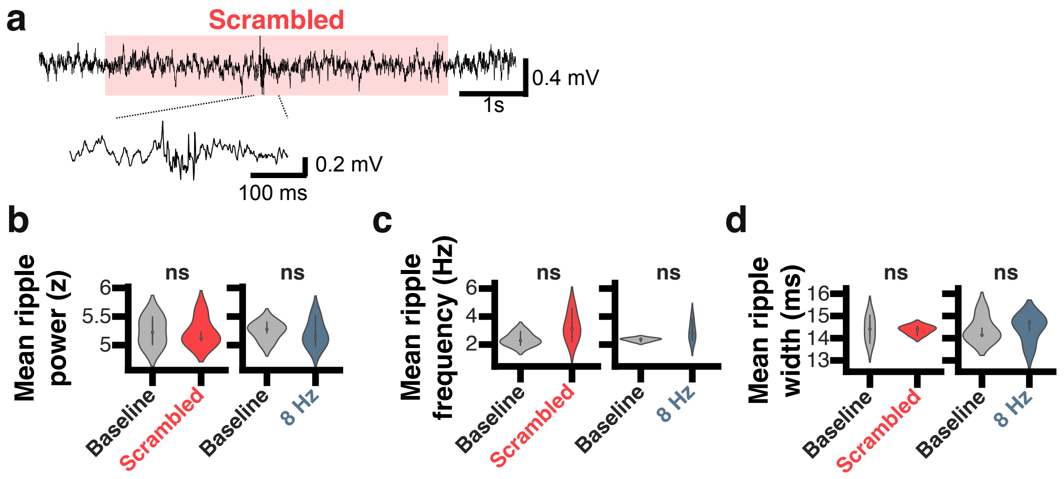


**Supplemental figure 4. Characteristics of sharp wave ripples are not affected by optogenetic stimulations. a,** example unfiltered trace of a ripple event during a scrambled optogenetic stimulation epoch. **b**, mean z-scored ripple power during baseline and either 8 Hz (blue) or scrambled (red) stimulations. **c**, same for mean ripple frequency of occurrence. **d**, mean ripple width.

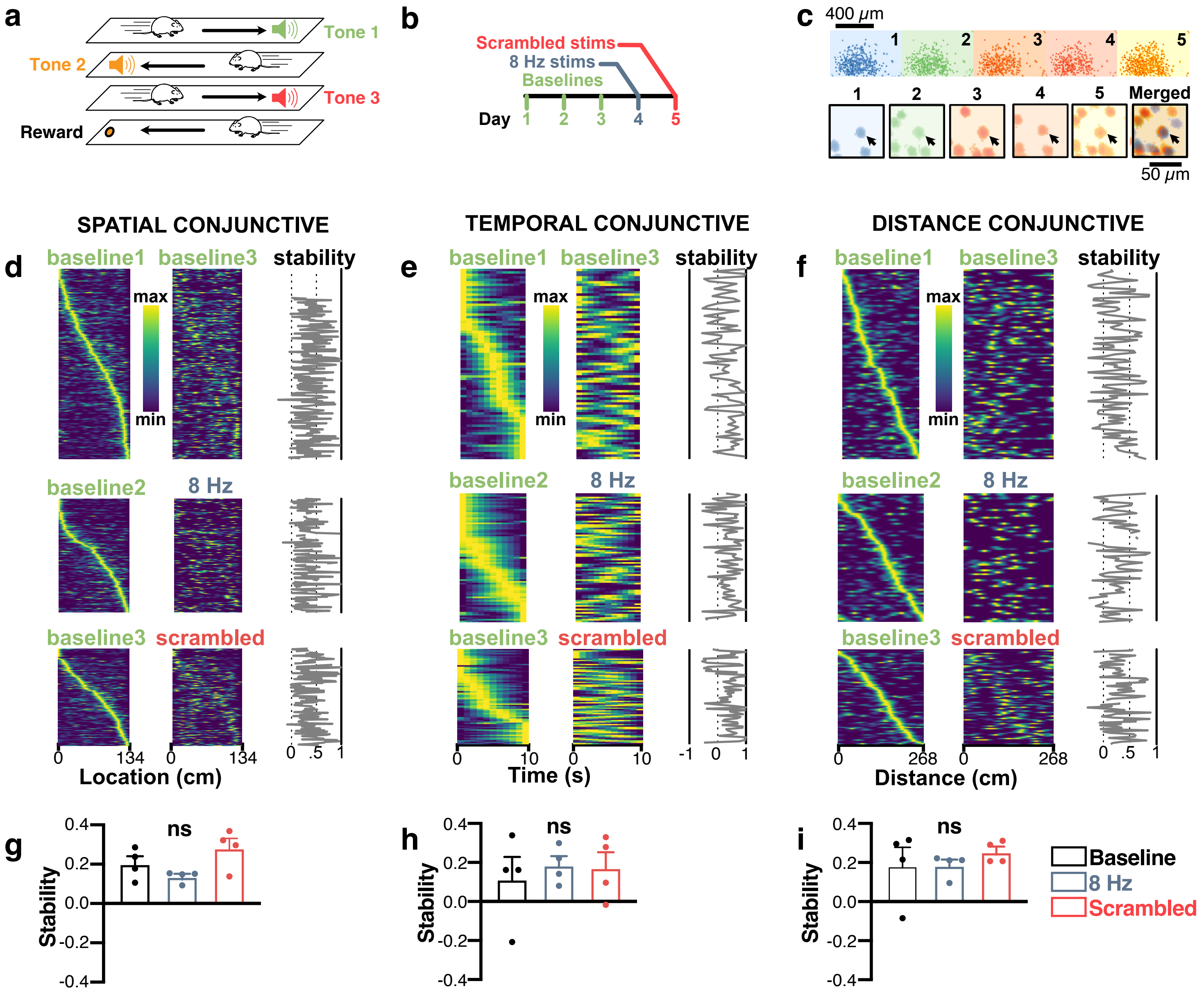


**Supplemental figure 5: MS optogenetic stimulation does not disrupt the stability of spatiotemporal codes in conjunctive cells. a**, mice were recorded on the 3-tone linear track to identify time-, place- and distance-modulated cells. **b**, experimental timeline. **c**, spatial footprints of neurons recorded over days and comparison scheme used to assess stability (top). Example neuron (black arrow) tracked over the 5 experimental days (bottom). **d**, sorted spatial tuning curves for identified conjunctive, spatially-modulated cells across day pairs. The first day is used as a baseline and mice undergo 8 Hz, scrambled, or no stimulations (control treatment) during the other day. Stability is computed as the pairwise correlation of fields between the two-day pairs. **e**, same for conjunctive, time-modulated cells. **f**, same for conjunctive, distance-modulated cells. **g**, corresponding average stability for conjunctive, spatially-modulated cells (each dot represents the average stability value for an individual mouse). **h**, same for conjunctive, time-modulated cells. **i**, same for conjunctive, distance-modulated cells.

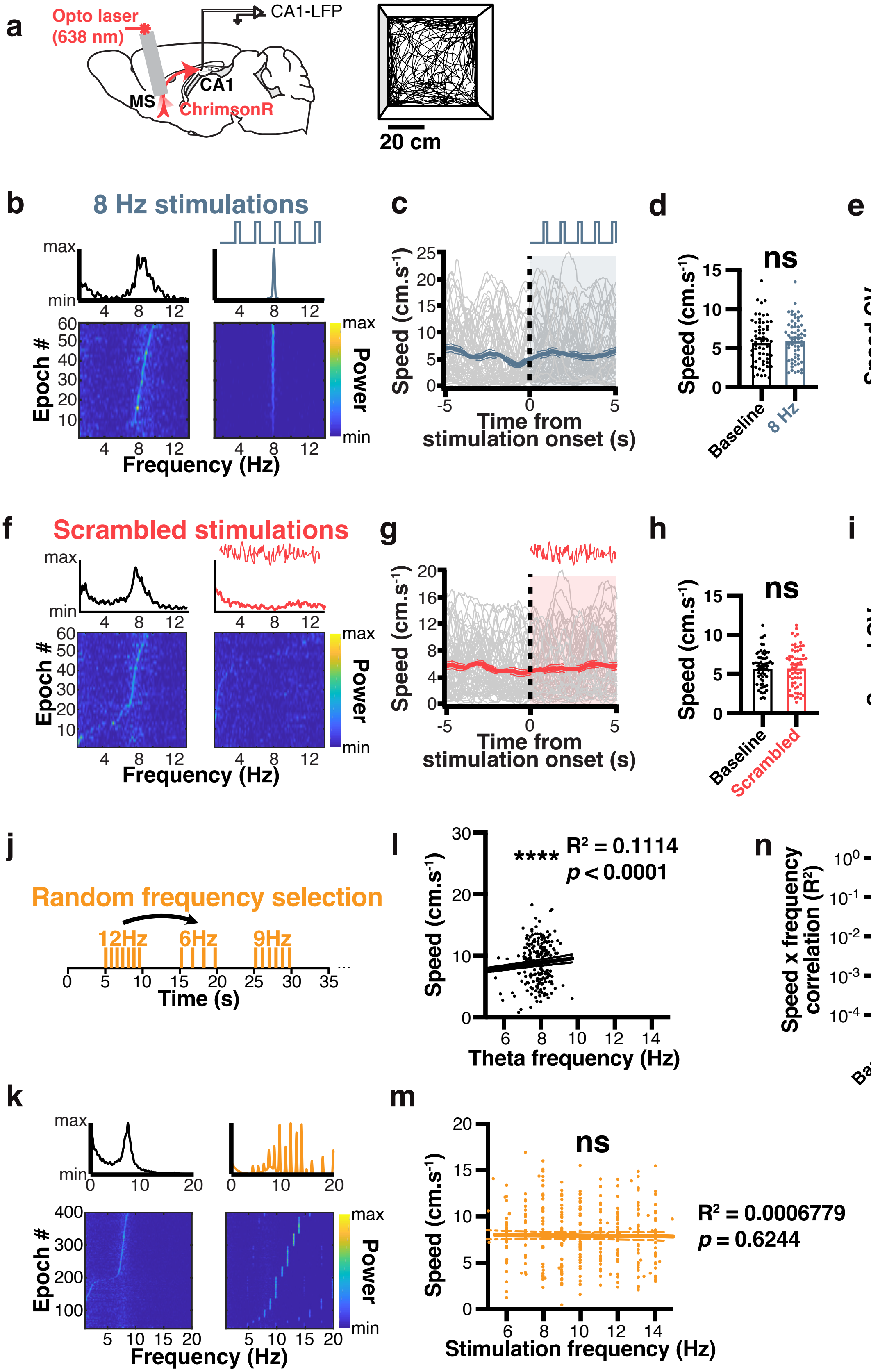

a
Opto laser (638 nm)
CA1-LFP
CA1
MS
ChrimsonR
20 cm
b
8 Hz stimulations
max
min
Epoch #
Frequency (Hz)
Power
c
Speed (cm.s-1)
Time from stimulation onset (s)
d
ns
Speed (cm.s-1)
Baseline
8 Hz
e
ns
Speed CV
Baseline
8 Hz
f
Scrambled stimulations
Epoch #
Frequency (Hz)
Power
g
Speed (cm.s-1)
Time from stimulation onset (s)
h
ns
Speed (cm.s-1)
Baseline
Scrambled
i
ns
Speed CV
Baseline
Scrambled
j
Random frequency selection
12Hz
6Hz
9Hz
Time (s)
l
****
R2 = 0.1114
p < 0.0001
Speed (cm.s-1)
Theta frequency (Hz)
n
Speed x frequency correlation (R2)
*
Baseline
Stimulation
k
Epoch #
Frequency (Hz)
Power
m
ns
Speed (cm.s-1)
R2 = 0.0006779
p = 0.6244
Stimulation frequency (Hz)

**Supplemental figure 6. MS optogenetic stimulation does not control locomotor speed. a,** mice were implanted with fiber optics in the MS after transfecting ChrimsonR, implanted with an electrode in CA1 (left), and freely explored an open field (right). **b**, average Fourier power spectra for 60 x 5s stimulation epochs (top), and sorted power spectra for each stimulation epoch (bottom) for baseline (left, in black), and 8 Hz (right, in blue) stimulations in one example mouse. **c**, peristimulus time plot of locomotor speed before and during 8 Hz MS optogenetic stimulation. Average speed in blue (line thickness indicated SEM). **d**, average locomotor speed before and during 8 Hz MS optogenetic stimulation. **e**, locomotor speed coefficient of variation (CV) before and during stimulation. **f**, average Fourier power spectra for 60 x 5s stimulation epochs (top), and sorted power spectra for each stimulation epoch (bottom) for baseline (left, in black), and scrambled (right, in red) stimulations in one example mouse. **g**, peristimulus time plot of locomotor speed before and during scrambled MS optogenetic stimulation. Average speed in red (line thickness indicated SEM). **h**, average locomotor speed before and during scrambled MS optogenetic stimulation. **i**, locomotor speed coefficient of variation (CV) before and during stimulation. **j**, random frequency generator used to pace theta oscillations at varying frequencies (see Methods). **k**, average Fourier power spectra for 400 x 5s stimulation epochs (top), and sorted power spectra for each stimulation epoch (bottom) for baseline (left, in black), and randomly selected (right, in orange) stimulations in one example mouse. **l**, relationship between locomotor speed and natural (unstimulated) theta frequency. **m**, same but during pacing of theta oscillations at varying stimulation frequencies. **n**, correlation between speed and natural (black) or controlled (orange) theta oscillations in N = 4 mice.

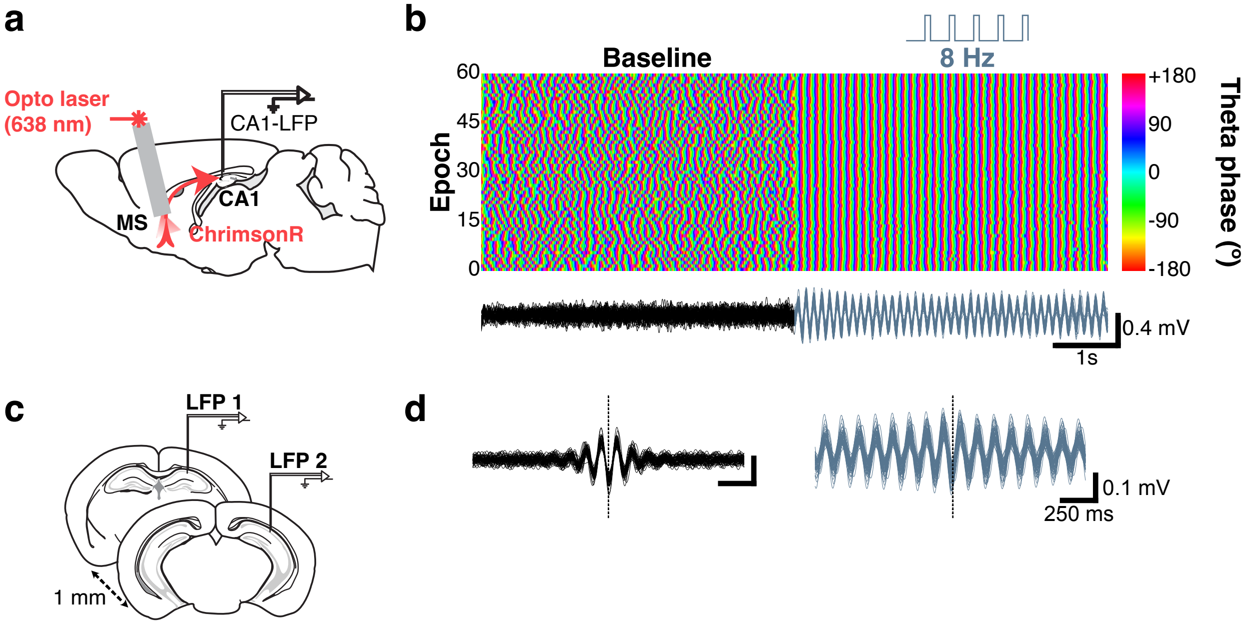


**Supplemental fig. 7. 8 Hz optogenetic stimulations lead to non-physiological theta synchrony. a**, mice were injected with ChrimsonR and implanted with a fiber optic in the MS. **b**, theta phase over time before and during optogenetic stimulation for (each horizontal line represents one of 60 epochs). Bottom, corresponding traces. **c**, mice were implanted with two electrodes in dorsal CA1, 1 mm in the septotemporal axis. **d**, cross-correlation between the two recording sites (LFP1 and LFP2 in **c**) before (black traces) or during 8 Hz optogenetic stimulations.
